# A CRISPR/Cas12a-assisted platform for identification and quantification of single CpG methylation sites

**DOI:** 10.1101/2021.04.06.438612

**Authors:** J.E. van Dongen, J.T.W. Berendsen, J.C.T. Eijkel, L.I. Segerink

## Abstract

Clustered Regularly Interspaced Short Palindromic Repeats (CRISPR)/associated nuclease (Cas) systems have repeatedly shown to have excellent performance in nucleotide sensing applications^1–5^. High specificity and selectivity of Cas effector proteins is determined by the CRISPR RNA’s (crRNA’s) interchangeable spacer sequence, as well as position and number of mismatches between target sequence and the crRNA sequence^1^. Some diseases are characterized by epigenetic alterations rather than nucleotide changes, and are therefore unsuitable for CRISPR-assisted sensing methods. Here we demonstrate a method to discriminate single CpG site methylation in DNA, which is an epigenetic alteration, by the use of methylation-sensitive restriction enzymes (MSREs) followed by Cas12a-assisted sensing. Non-methylated sequences are digested by MSREs, resulting in fragmentation of the target sequence that influences the R-loop formation between crRNA and target DNA. We show that fragment size, fragmentation position and number of fragments influence the subsequent collateral *trans*-cleavage activity towards single stranded DNA (ssDNA), enabling deducting the methylation position from the cleavage activity. Utilizing MSREs in combination with Cas12a, single CpG site methylation levels of a cancer gene were for the first time determined. The modularity of both Cas12a and MSREs provide a high level of versatility to the Cas12a–MSRE combined sensing method, which opens the possibility to easily and rapidly study single CpG methylation sites for disease detection.

---

CRISPR/Cas systems have revolutionized biotechnology and synthetic biology^6–9^. Of special interest for the sensor field are the type V and VI effector proteins, two subclasses of the class 2 Cas proteins. These proteins showcase, besides targeted *cis-*cleavage, untargeted *trans*-cleavage activity. This *trans*-cleavage activity has multiple turnovers, enabling a signal amplification of up to 10,000^5^, which can be deployed in sensor applications to lower the limit of detection.

For double stranded DNA (dsDNA) sensing, type V effector protein Cas12a is mostly used. Critical for both *trans-* and *cis-*cleavage activities of Cas12a is the R-loop formation step, where the crRNA and target strand of the dsDNA hybridize and the catalytic site is revealed (Extended Data Fig. 1). Due to the late transition state for R-loop formation, Cas12a is generally able to discriminate against mismatches across the R-loop, determining its low mismatch tolerance^10^. However, the degree of mismatch tolerance greatly depends on the origin of the effector protein and the position of the mismatch^1^. The nucleotides in the so-called ‘seed’ region hybridize with the target protospacer adjacent motif (PAM) proximal nucleotides and are considered critical in target affinity^11^, and mismatches in this region strongly negatively affect the *trans-*cleavage activity. PAM-distal mismatches show a much lower effect on the *trans-*cleavage activity, and only 15 out of 23 PAM-proximal matching nucleotides are needed for (temporary) binding of (enzymatically deactivated) Cas12a^12^.

While many diseases could be related to single changes in the DNA nucleotides, not all genetic modifications related to diseases include changes in nucleotide sequences. In the last decade, cancer biomarker research has focused on biomarkers representing epigenetic alterations associated with the development of cancer^13^. One of these epigenetic events includes DNA (hyper)methylation. Methylated cytosines in CpG dinucleotides are an epigenetic alteration to the DNA, that is involved in both healthy processes like embryogenic development and transcriptional regulation, as well as in less benign processes such as autoimmune diseases and several cancer types^14–16^. These alterations can generally not be differentiated by Cas12 as a mismatch and are therefore, in principle, unsuitable for CRISPR sensing methods. Li *et al*. bypassed this issue by combining their HOLMESV2 with bisulfite conversion. However, bisulfite-based techniques are time consuming and labor-intensive chemical treatments that damage DNA, induce (unwanted) DNA fragmentation, have limited throughput, and dramatically reduce the genome complexity, making specific targeting of sequences more difficult^17^. An alternative method to bisulfite conversion is the use of MSREs. Conventionally, MSREs are combined with quantitative polymerase chain reaction (qPCR) amplification for quantification purposes. However, this comes with several drawbacks: qPCR requires relatively long amplicon lengths (70-200 bp), and these amplicons should contain at least two MSRE restriction sites to reliably inhibit amplification for non-methylated samples^18^. Therefore, it is not possible to investigate the methylation levels of single CpG dinucleotides, which has been shown to be strongly associated with the development of gastric, liver and colon cancers^19,20^.

A method to bypass both the long amplicon length and the need of multiple restriction sites to visualize the methylation level could be MSRE-based CRISPR/Cas sensing. In this case of restriction by MSREs prior to addition of the Cas effector protein, the fragmentation would affect the R-loop formation, thereby slowing down the *trans-*cleavage activity.

## Shorter target fragments decrease, and longer target fragments increase *trans*-cleavage activity

When combining MSREs with Cas12a, the R-loop formation between the crRNA and non-methylated target DNA strands will be disrupted. To our knowledge the effect of the resulting short(er) target fragments complementary to crRNA was only previously touched upon by Chen *et al*., who showed that target strand cleavage by Cas12a is not required to trigger the non-specific collateral cleavage activity towards ssDNA, but decreases the amount of *trans-*cleavage activity dramatically^1^. From previous work by Swarts and Jinek^21^ and Chen *et al*.^1^ we hypothesize that the most important prerequisite for *trans*-cleavage is the structural rearrangement of Cas12a to reveal the RuvC catalytic site by the R-loop formation, followed by clearance (See Supplementary information, section 1). This clearance can result either from *cis*-cleavage of target strands followed by diffusion of the PAM-distal fragment, or is not needed, due to the binding of short(er) target strands that do not interfere with the RuvC catalytic site. Therefore, we expect a tradeoff between R-loop formation length and clearance of the RuvC catalytic site.

To experimentally quantify the effect of different target sequence lengths on Cas12a *trans-*cleavage, we tested the ability of synthetic oligonucleotides of different lengths to activate the *trans*-cleavage activity of Cas12a. Different target sequences were added to Cas12a and *trans*-cleavage was followed resulting in an increase in fluorescence by addition of a reporter DNA, which contained a fluorophore-quencher pair. To see whether the GC-content of the target sequences influenced the activity, we selected the MAL gene with a high GC-content, for which it is known that its methylation is related to several cancer types^22–26^, and a randomly designed sequence with a low GC-content (Extended Data Fig. 2). Besides the GC-content, the spacer length of the crRNA was varied as well, as it is known from literature that this length could strongly affect the cleavage efficiency of Cas12a^27,28^. Figure 1a and 1b show the results of different target fragment lengths of the high GC-content MAL gene at different concentrations (for 0.1 nM see Extended Data Fig. 3). Independent of the GC-content or crRNA length, the highest *trans*-cleavage activity could be observed for fragments from 15 bp match to the crRNA and onwards, which is in agreement with the length of PAM-distal mismatches that are accepted for enzymatically deactivated Cas12a binding^12^. Remarkably, the highest *trans*-cleavage activity of Cas12a is not observed for a DNA fragment that spans the total crRNA length, but for shorter fragments. This could be explained by a conserved aromatic residue (W382) of Cas12a that stacks on position 20 of the crRNA, preventing further R-loop propagation of the RNA:DNA duplex ^10,29–31^ and causing bases on position 20 and onwards to not further contribute to revealing RuvC for ssDNA access for *trans*-cleavage activity. Since we observe a slower *trans*-cleavage rate for longer fragments, we propose that steric hindrance by the staggered cut-end of the target DNA strand could induce a slower *trans*-cleavage activity. Shorter target lengths do not interfere with the RuvC catalytic site, and therefore enable direct *trans*-cleavage of ssDNA, without dependence on diffusive clearance or steric hindrance of the (remaining) *cis-*cleavage product. This also supports the observation that a decreased spacer length of the crRNA increases the *trans*-cleavage for both low GC containing and high GC containing sequences, which originates probably also from steric hindrance of the longer spacer sequence.

**Figure 1.**
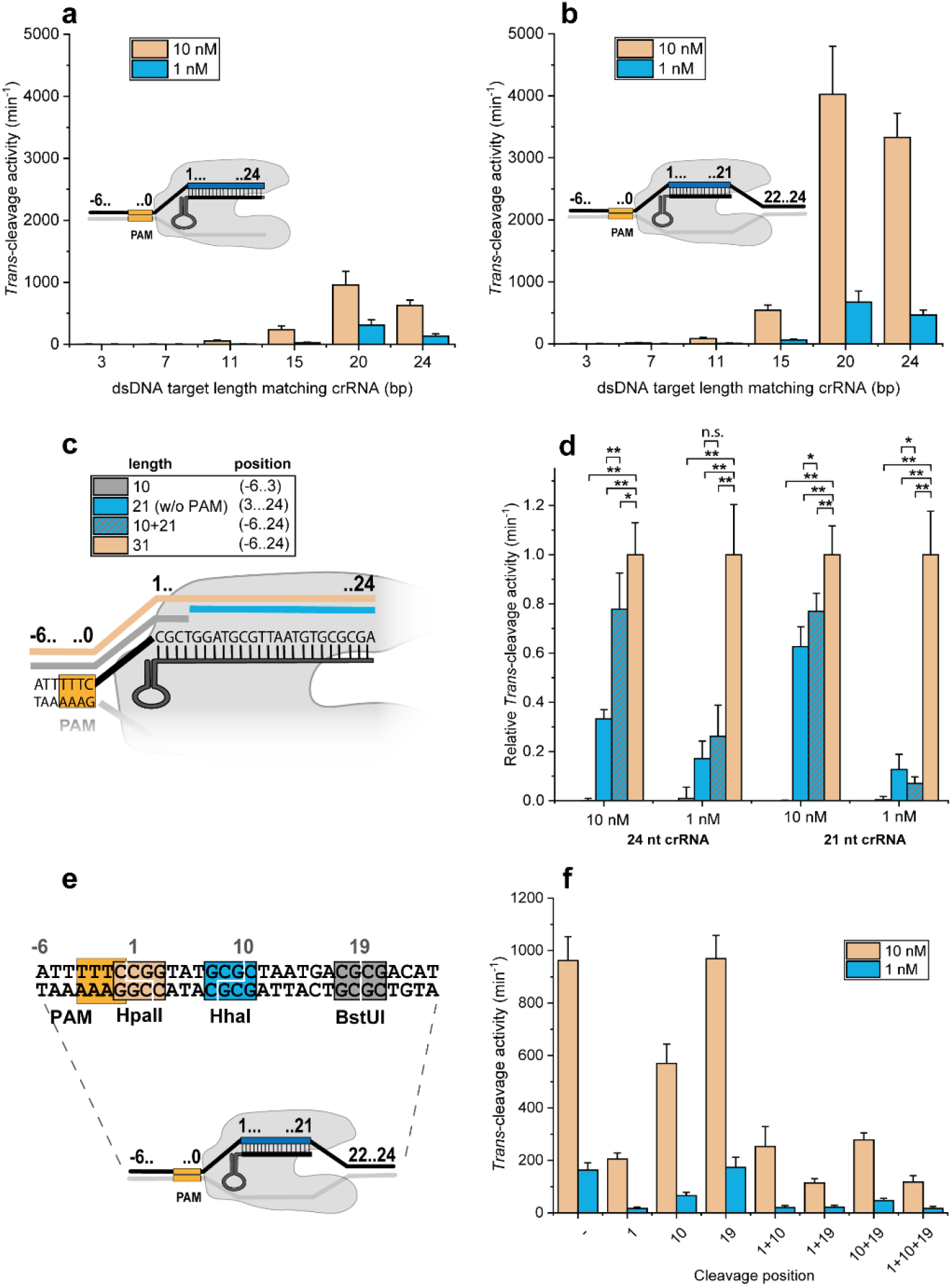
Quantification of *trans-*cleavage activity using different dsDNA target fragments. High GC-content MAL sequence with a 3, 7, 11, 15, 20 and 24 bp target sequence that matches with a Cas12a ribonucleotide complex with a) a 24 nt crRNA spacer length and b) a 21 nt crRNA spacer length. c) Cartoon representation of Cas12a ribonucleotide complex with a 24 nt crRNA complex that binds to 10, 21, 10+21 and 31 bp fragments. d) Relative *trans-*cleavage rates of these different target lengths with the 24 and 21 nt spacer-crRNA. ** indicates p<0.001 and * indicates p<0.05 (Two-sample T-test). e) Cartoon representation of the cleavage position of the MSREs on a 31 bp target sequence related to the crRNA. f) *Trans*-cleavage activity of a 31 bp total dsDNA target sequence, after treatment with a single type or a mixture of MSREs. In all figures error bars represent the mean +/- s.d., where n=9 (three replicates for three independent targets).

## Increased *trans-*cleavage in the presence of two fragments both complementing the crRNA spacer

Whereas experiments with different fragment lengths gave us some insight in the effect of shorter fragment lengths on the Cas12a *trans*-cleavage activity, utilizing MSREs will result in all fragments being present in the mixture in equimolar concentrations. In theory, this allows all fragments to bind either individually or collectively to the crRNA. To mimic this situation, we caried out experiments with two fragments (21 bp and 10 bp) that together span the total 31 bp target sequence under the same experimental conditions as the fragmented targets (Figure 1c). Figure 1d shows the *trans-*cleavage activity of the different fragment lengths and the equimolar combination of the 10 and 21 bp fragments normalized to the *trans*-cleavage activity of 31 bp fragments at the same concentration. Remarkably, the activity resulting from 10 + 21 bp is in most conditions significantly higher than the sum of the activities for the individual 10 and 21 bp fragments (p<0.05), suggesting that both fragments can bind simultaneously to the crRNA, inducing a larger R-loop formation, resulting in higher *trans-*cleavage activity. The same trend was observed for low GC-content target sequences (Extended Data Fig. 4), although the contribution of both fragments being present at equimolar concentrations was percentual less compared to high GC containing sequences. Additional experiments with high GC-content MAL sequences confirm that both fragments bind to the crRNA simultaneously (Extended Data Fig. 5). However, the simultaneous binding of both fragments decreases significantly at lower target DNA concentrations (p<0.001 for both crRNA lengths), which is observed as a relatively lower contribution of both fragments to the *trans*-cleavage activity for both high GC as low GC containing sequences.

## T*rans*-cleavage activity: cleavage position and fragment number matters

We next tested whether the cleavage position on the target-DNA relatively to the PAM would affect the *trans*-cleavage activity. We designed a 31 bp dsDNA sequence with three MSRE recognition sites, which allowed us to selectively induce cleavage at different positions (Figure 1e). After overnight digestion with one single type or cocktails of MSREs, the Cas12a with associated crRNA was added and *trans*-cleavage was followed. Figure 1f shows the results of the *trans-*cleavage activity. PAM-proximal cleavage sites show the largest effect on the *trans-*cleavage activity, while PAM-distal cleavage at position 19 has no significant influence, which is in accordance with results discussed in previous work^10,29–31^. Combining cleavage sites decreases the relative *trans*-cleavage activity. This experiment gives crucial information for the crRNA design in single site CpG experiments, since the highest sensitivity is achieved for PAM-proximal non-methylated cytosines that are targeted by MSREs.

## Single CpG site methylation level quantification by combining MSREs and Cas12a sensing

With the previous experiments we have confirmed that target DNA fragments without a PAM-proximal sequence result in a significantly lower *trans*-cleavage activity of Cas12a compared to sequences that match the entire crRNA sequence, or miss PAM-distal fragments even when both fragments are still present in the same solution. This opens the door for single CpG site methylation analysis, utilizing MSREs for fragmentation purposes. For these experiments, longer MAL gene DNA fragments of 120 bp were used to mimic a more realistic DNA fragment, as can typically be found in liquid biopsies like urine and blood^32–34^. We selected a crRNA that targets a single, PAM-proximal CpG site to achieve the highest sensitivity, as discussed in the previous section.

Figure 2a shows a schematic of the experimental outline for the detection of the methylation percentage of a single CpG site in the MAL gene. Since synthetic oligonucleotides are used in these experiments, methylation was performed with the M.SssI enzyme that, in principle, should methylate all CpGs of the oligonucleotide. Mock digestions were performed so that methylated and non-methylated DNA could be mixed in different ratios, to mimic different methylation percentages. Discrimination between methylated and non-methylated DNA is performed by AciI, which is a MSRE, with a recognition site that is PAM-proximal to the crRNA selected and results in two fragments that can be compared to the 10+21 bp fragments tested in the previous section. Using gel electrophoresis for fragmentation monitoring (Figure 2b), we observed some cleavage for the M.SssI-treated sample in the presence of AciI which suggests that either the M.SssI methylation was not complete, or AciI poses some non-specific activity resulting in underestimation of the methylation percentage. However, since we performed a calibration step this is not an issue for these experiments. Depending on the application and clinical setting of the assay one could optimize the MSRE digestion, resulting in either more false positives or false negatives.

**Figure 2.**
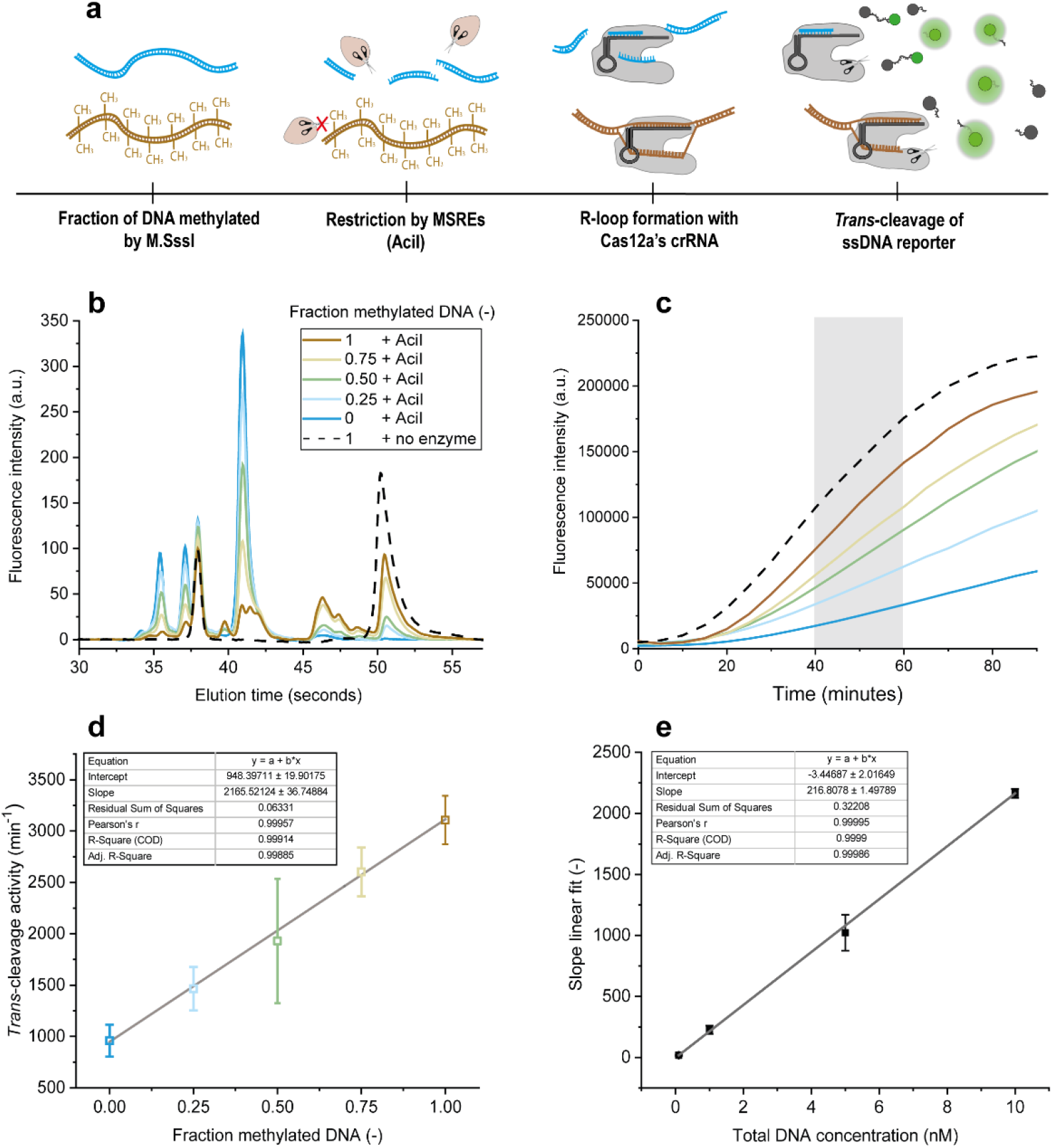
a) Schematic of time line for detection of methylation percentage of a single CpG site with MSREs and Cas12a. b) gel electrophoresis of methylated DNA without enzyme treatment and different fractions of methylated : non-methylated DNA after enzyme treatment. c) Increase in fluorescence over time for different fractions of methylated : non-methylated DNA after enzyme treatment at a total DNA concentration of 10 nM, in gray the time slot used for determination of the *trans-*cleavage activity. d) *Trans-*cleavage activity of different methylated DNA : non methylated DNA fraction at a total DNA concentration of 10 nM. Error bars represent the mean +/- s.d., where n=9 (three replicates for three independent targets). e) slope of the linear fit for the relation between fractions of methylated : non-methylated DNA against the *trans-*cleavage for different total DNA concentrations (0.1, 1, 5 and 10 nM). Error bars represent the mean +/- s.d. of the linear fit through the 5 data points per concentration.

After MSRE treatment, the different ratios methylated : non-methylated DNA were diluted to total DNA concentrations of 10, 5, 1 and 0.1 nM and added to Cas12a with a crRNA spacer length of either 21 or 24 nt. Figure 2c shows the increase in fluorescence for the different ratios at a total DNA concentration of 10 nM with a crRNA length of 21 nt. The average slope of the linear section of these curves between 40 and 60 minutes was used to determine the average *trans-*cleavage activity. The effect of methylation on the *trans*-cleavage activity of Cas12a could be neglected, as this did not result in significant differences in average *trans*-cleavage activity (Extended Data Fig. 6). Figure 2d shows a linear fit with a R^2^ =0.999 when plotting this *trans-*cleavage activity against the fraction of methylated DNA (for all other concentrations see extended Data Fig.7 for 21 nt crRNA-spacer). The slopes of the curves corresponding to all different total DNA concentrations were plotted against the total DNA concentration in Figure 2e, additionally showing the excellent sensitivity of Cas12a towards different DNA concentrations. Utilizing 24 nt crRNA containing Cas12a resulted in similar linearity over both the fraction methylated DNA and the total DNA concentration (extended Data Fig. 8).

## Discussion

The methylation quantification method presented here for combined Cas12a–MSRE sensing opens the possibility to identify individual CpG methylation sites in the genome by customizing the spacer sequence, which is the section of the crRNA that hybridizes with the target DNA strand. The typical length of this section of 20-24 nt offers the possibility of targeting unique sequences in most living species. For example, in the human genome, 60% can be targeted as a unique sequence by customizing the crRNA sequence^35^. Limitation of Cas12a is the PAM sequence, however, the world of biotechnology is making great process in the design of a PAM-free Cas nuclease, which will increase the possibilities to target sequences tremendously^36^. Without the need of a PAM recognition sequence and the information given in this paper the best possible crRNA-spacer sequences could be designed that offers the highest sensitivity for targeting single CpG sites in the entire genome.

This new approach to epigenetic sequencing will allow extracting methylation patterns, providing as yet unknown patient data with the potential to revolutionize disease diagnostics. Furthermore, restriction enzymes can be exploited that can discriminate 5-methylcytosine (5-mC) from 5-hydroxymethylcytosine (5-hmC), which is one of the hot-topics in the epigenetics field^37^. The revolution in biotechnology also resulted in many manufactures providing ultra-fast enzymes that offer qualified digestion in 5-15 minutes. In combination with the relatively fast CRISPR/Cas12a based fluorescence detection (about 15 min − 1 hour depending on the input concentration), this method could easily be adapted to a point-of-care method, allowing quantitative and sensitive analysis in resource-poor settings^38^.

## Acknowledgements

We would like to thank The Weijerhorst team from the University of Twente and the AmsterdamUMC for the fruitful collaboration. The Weijerhorst Foundation is acknowledged for their financial support.

