## Supplementary material for "A CRISPR/Cas12a-assisted platform for identification and quantification of single CpG methylation sites": Methods

### Oligonucleotides

All DNA oligonucleotides were synthesized by Eurofins Genomics and the sequences are available in the supplementary Table 1. All crRNA fragments were Alt-R® A.s. Cas12a crRNAs, synthesized by Integrated DNA Technologies (IDT, Coralville IA). All sequences can be found in the supplementary information. The PAM sequence is underlined and the sequence that complements to the crRNA is in bold.

### Cas12a ribonucleoprotein complex formation

ALT-R CRISPR-Cas12a (Cpf1) Ultra (Integrated DNA technologies) and custom crRNA were mixed in a ratio 1:2 for 30 minutes at room temperature to form ribonucleotide protein (RNP) complexes. The assembled complex was then 100 times diluted in RNAse free water (MACHEREY-NAGEL) and stored in the freezer at -20°C until further use.

### Cas12a collateral cleavage assay

In a total volume of 70 µl, 1x NEBuffer 2.1 (New England Biolabs), 20 nM of RNP complex, 1000 nM of fluorophore-Quencher “reporter” DNA (Eurofins Genomics) were mixed with different concentrations of target dsDNA. Mass screening experiments were performed in a black polypropylene 384 well plate (Corning®), where each well was filled with 20 µl of reaction mixture, allowing triplicates of the total reaction volume. In these mass screening experiments, a MasterMix was prepared containing the RNP, NEB2.1 and reporter DNA. Both the MasterMix and reaction mixture were pipetted on ice and the reaction mixture transferred to a frozen, black 384-well plate to inhibit reactions until the plate was transferred to the plate reader.

### *Trans*-cleavage activity determination of short dsDNA fragments

To study the effect of different fragment sizes on the *trans*-cleavage activity of Cas12a, two synthetic 31 base pair fragments (Eurofins Genomics) were selected with different methylation levels. Besides the full-length sequence, different dsDNA sequence lengths were tested (10, 14, 18, 22, 27 bp) that span from 3 nucleotides before the PAM sequence until the length of interest (Eurofins Genomics). RNPs were formed as described in a previous section and included both 24 and 21 nt spacer length containing crRNAs (Integrated DNA technologies). The increase in fluorescence was followed in a mass-screening experiment, as described in the previous section. *Trans*-cleavage was determined by taking the first derivative of the fluorescence increase curves and averaging the *trans*-cleavage over 5 measurement points (25 minutes) in the linear range of the fluorescence-increase curve.

### *Trans*-cleavage activity determination of short dsDNA fragments combined

To study the effect of different fragment sizes present in the same molar ratios on the *trans*-cleavage activity of Cas12a, two synthetic base pair fragments (Eurofins Genomics) of a total length of 10 and 21 bp were selected. Experiments as described in the previous section were performed (with the one exception that both fragments were added at the same concentration) and experiments were the fragments were incubated with the RNP for ½ hour or 1 hour without the reporter ssDNA. For the 10 and 21 bp fragments, four different experimental conditions were tested, where either 10 or 21 bp fragments were incubated for ½ hour prior addition of the 21 or 10 bp fragment followed by ½ hour of incubation, as well as immediately addition of both fragments and incubation of either ½ hour or 1 hour. After the total incubation time (max 1 hour) the RNP mixtures were placed on ice and the reporter ssDNA was added, before the plate was transferred to a multi-mode plate reader (BioTek) at room temperature, taking fluorescence measurements every 5 minutes (λex: 495; λem: 520). *Trans*-cleavage was determined by taking the first derivative of the fluorescence increase curves and averaging the *trans*-cleavage over 5 measurement points (25 minutes) in the linear range of the fluorescence-increase curve.

**Methylation sensitive restriction-enzyme for methylation "sequencing"**

In vitro methylation sequencing of 31 bp synthetic oligonucleotides was achieved by using a cocktail of different enzymes (each 5 U), consisting of HpaII, HhaI, SnaBI and/or BstUI (New England Biolabs). Mock digestions were performed under the same conditions, but without enzyme addition. DNA and enzymes were incubated in CutSmart Buffer (New England Biolabs), and reactions were performed overnight at 37°C or 60°C (depending on the enzyme), followed by an inactivation step at 85°C for 20 minutes in a T100 thermal cycler (BioRad). The digested DNA samples were analyzed for digestion by automated gel electrophoresis (ExperionTM, BioRad), and used as a target sequence in mass screening experiments to follow the corresponding Cas12a *trans*-cleavage activity. *Trans*-cleavage was determined by taking the first derivative of the fluorescence increase curves and averaging the *trans*-cleavage over 5 measurement points (25 minutes) in the linear range of the fluorescence-increase curve.

**Methylation sensitive restriction-enzyme digestion for Methylation % determination**

In vitro methylation of synthetic oligonucleotides was achieved in house, utilizing MSssI CpG-methyltransferase (New England Biolabs) according to manufacturer's protocol. For methylation-sensitive restriction enzyme experiments, both methylated and non-methylated DNA was mixed in different ratios as in input for digestion reactions with AciI (5 U: New England Biolabs R0551S). Mock digestion was performed under the same conditions, but with no added enzyme. DNA and enzymes were incubated in CutSmart Buffer (New England Biolabs), and reactions were performed overnight at 37°C in a T100 thermal cycler (BioRad), followed by an inactivation step at 65°C for 20 minutes. The digested DNA samples were analyzed for digestion by automated gel electrophoresis (ExperionTM, BioRad), and used as a target sequence in mass screening experiments to follow the corresponding Cas12a *trans*-cleavage activity as described in the previous sections. *Trans*-cleavage was determined by taking the first derivative of the fluorescence increase curves and averaging the *trans*-cleavage over 5 measurement points (25 minutes) in the linear range of the fluorescence-increase curve.

**Data analysis and statistics**

Data analysis and processing was carried out in OriginPro 2019. For statistical analysis two-sample t-test was performed to define significant differences between the *trans*-cleavage activity for different conditions.
