## Extended Data for "A CRISPR/Cas12a-assisted platform for identification and quantification of single CpG methylation sites"

*J.E. van Dongen[1]; J.T.W. Berendsen[1]; J.C.T. Eijkel[1]; L.I. Segerink[1]*
BIOS Lab on a Chip group, MESA+ Institute for Nanotechnology, Technical Medical Centre, Max Planck Institute for Complex Fluid Dynamics, University of Twente, Enschede, the Netherlands

### Extended Data

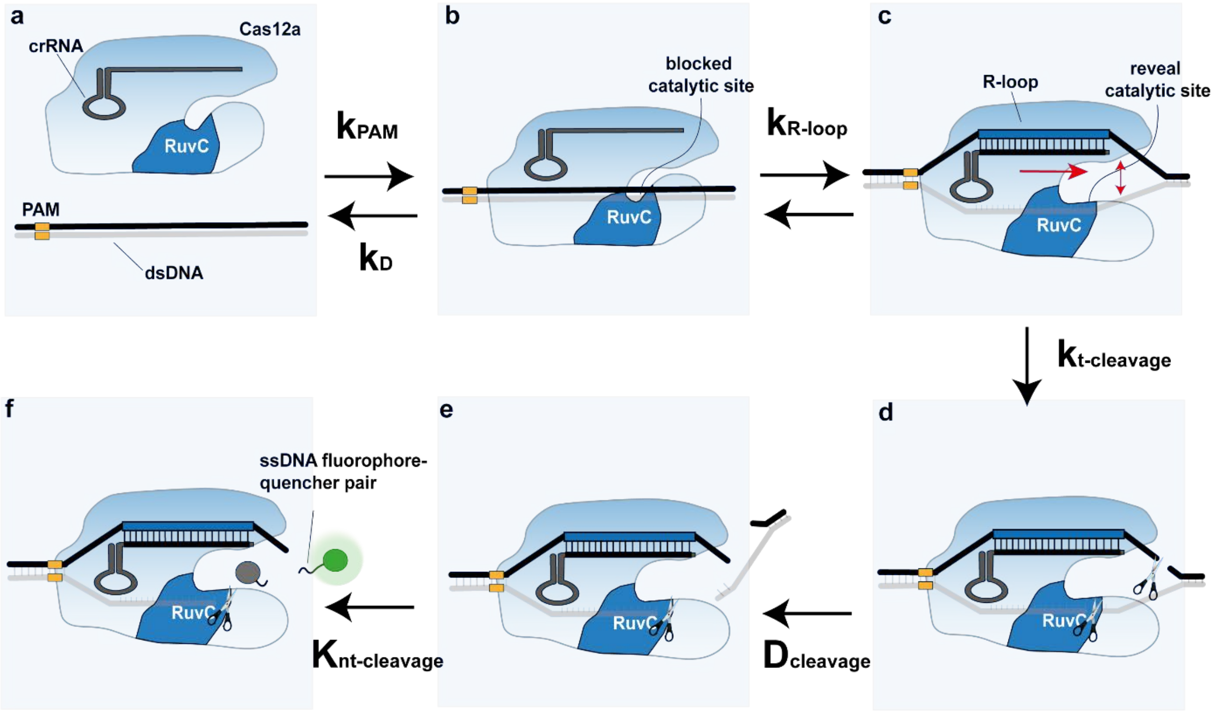


**Extended Data Fig. 1, schematic overview of the mechanism behind Cas12a's targeted *cis*- and non-targeted *trans*-cleavage. A) Cas12a and its crRNA are able to recognize dsDNA targets, which can be described by the association constant $K_{PAM}$, this binding is reversible and therefore the dissociation constant $K_D$ is included. B) Target recognition by Cas12a starts with PAM sequence recognition, promoting the unwinding of dsDNA. C) If part of the unwound dsDNA is complementary to the crRNA, R-loop formation takes place simultaneously with the unwinding of the dsDNA. This crRNA : DNA duplex formation starts from the PAM-distal nucleotides and promotes rearrangement of the Cas12a protein, revealing the RuvC catalytic site needed for both *cis*- and *trans*-cleavage. D) *Cis*-cleavage takes place for targeting DNA strand and its complementary strand, from this moment the binding reaction is in reversible. E) the PAM-distal sequence is released, while the PAM-proximal sequence remains bound to the Cas12a protein. The RuvC catalytic site remains blocked until the fragment diffused away from the catalytic site (determined by the diffusion coefficient $D_{cleavage}$). F) Cas12a remains catalytically active, allowing the *trans*-cleavage of non-targeted ssDNA strands, which can be followed by ssDNA-fluorophore quencher pairs, defined by the interacting constant of ssDNA to the Cas12a.**

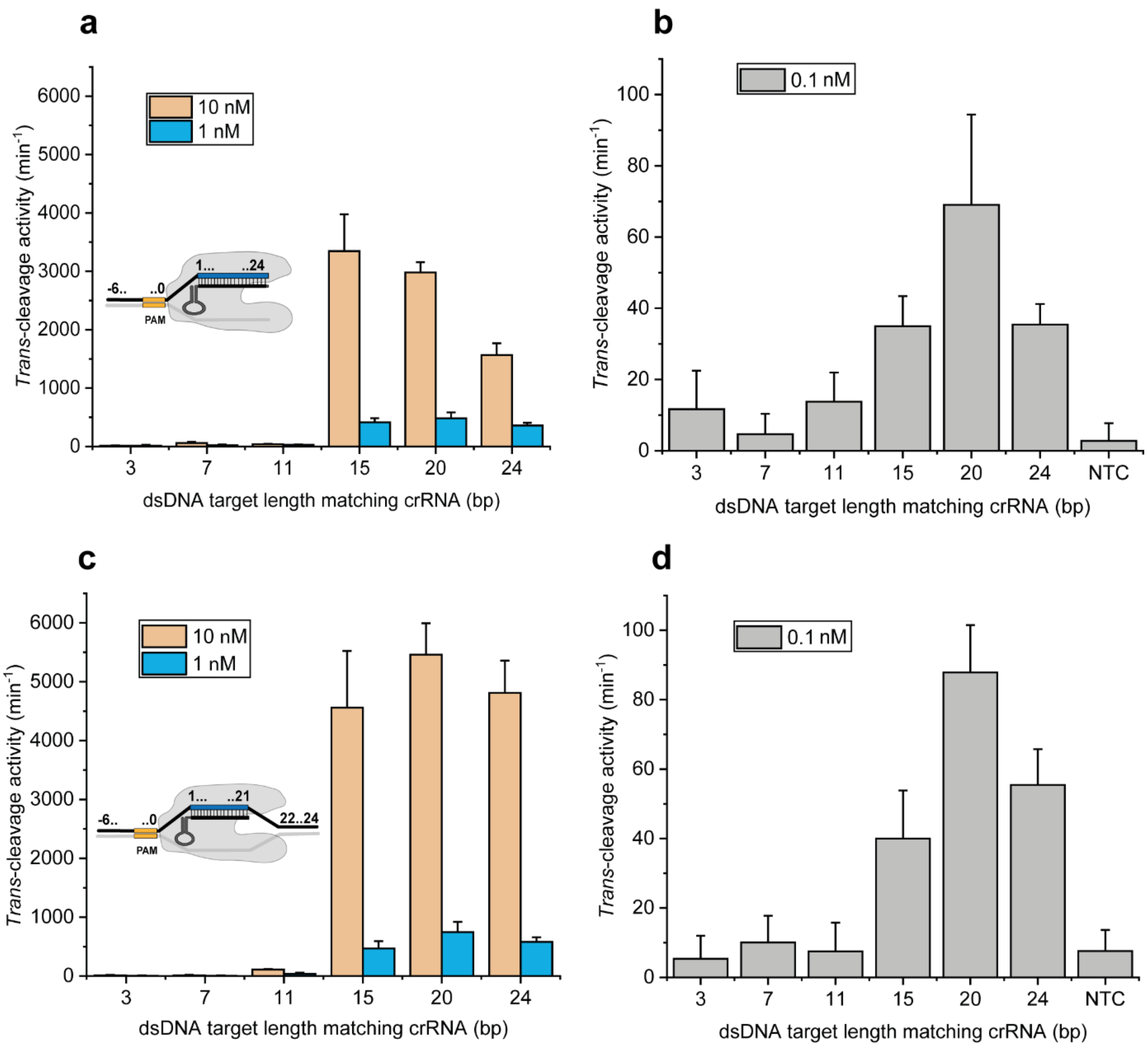


**Extended Data Fig. 2, Quantification of *trans*-cleavage activity using different dsDNA target fragments. Low GC-content sequence with a 3, 7, 11, 15, 20 and 24 bp target sequence that matches with a Cas12a ribonucleotide complex with a) a 24 nt crRNA spacer length at 10 and 1 nM and b) 0.1 nM and c) a 21 nt crRNA spacer length at 10, 1 nM and d) 0.1 nM. Error bars represent the mean +/- s.d., where n=6 (three replicates for two independent targets).**

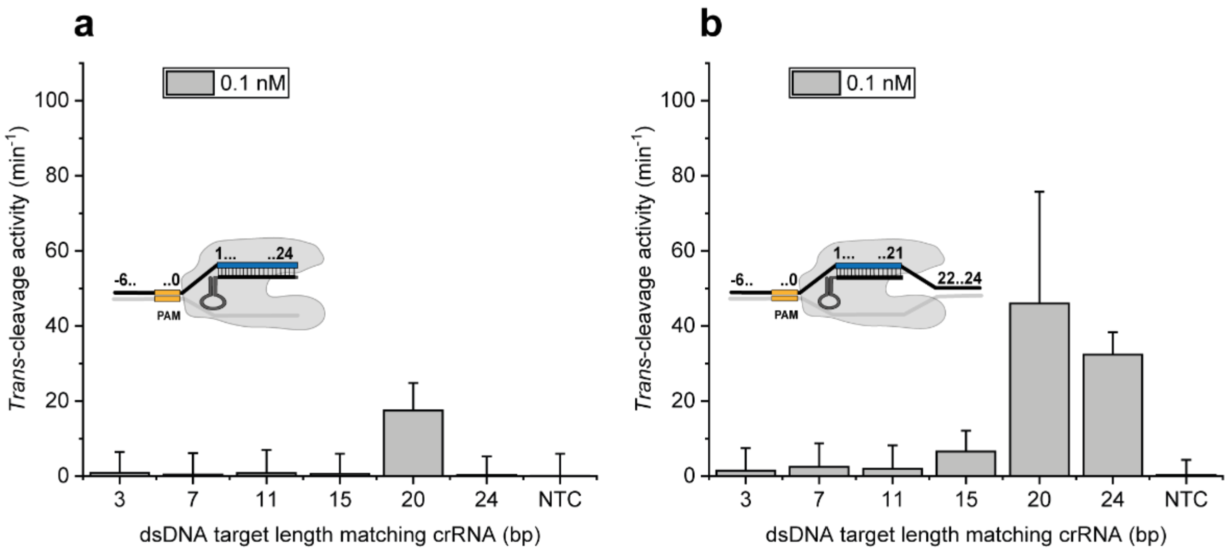


**Extended Data Fig. 3, Quantification of *trans*-cleavage activity using high GC-content MAL dsDNA target fragments. with a) a 24 nt crRNA spacer length and b) a 21 nt crRNA spacer length. Error bars represent the mean +/- s.d., where n=9 (three replicates for three independent targets).**

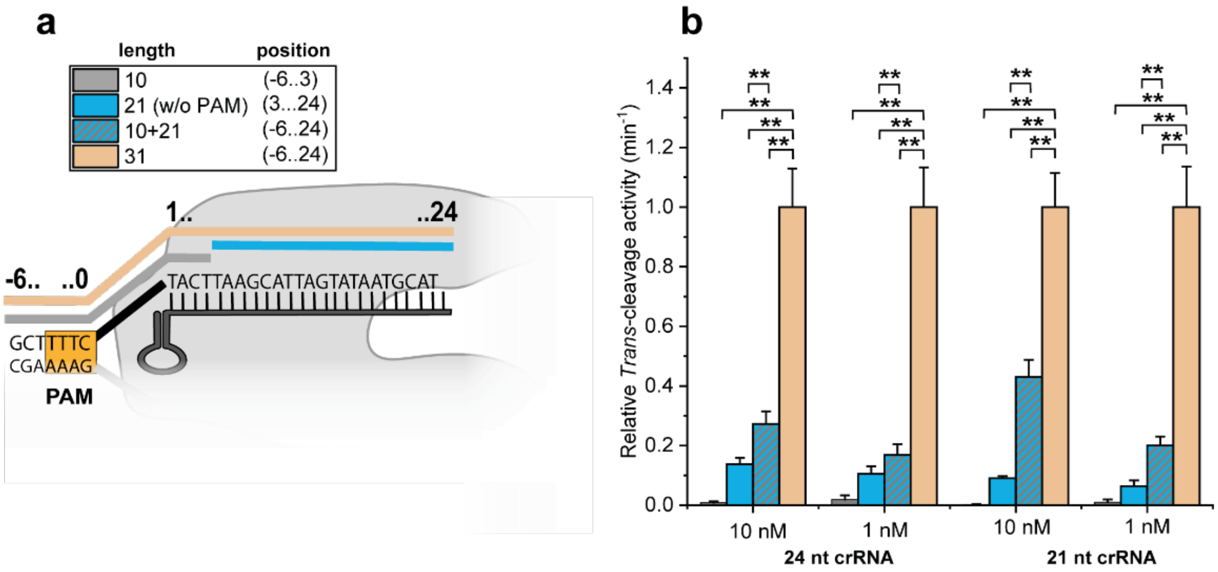


**Extended Data Fig. 4, Relative *trans*-cleavage rate of different fragment lengths for low GC-content fragments, normalized to the rate of 24 nt crRNA : target DNA match (31 bp total length) at the same concentration. Besides 10, 21 and 31 bp fragments, also 10 + 21 nt matches at equimolar ratios were included to mimic the situation after MSREs cleavage where fragments are present at similar ratios. ** indicates p<0.001. Experiments carried out with n=6 (three replicates for two independent targets).**

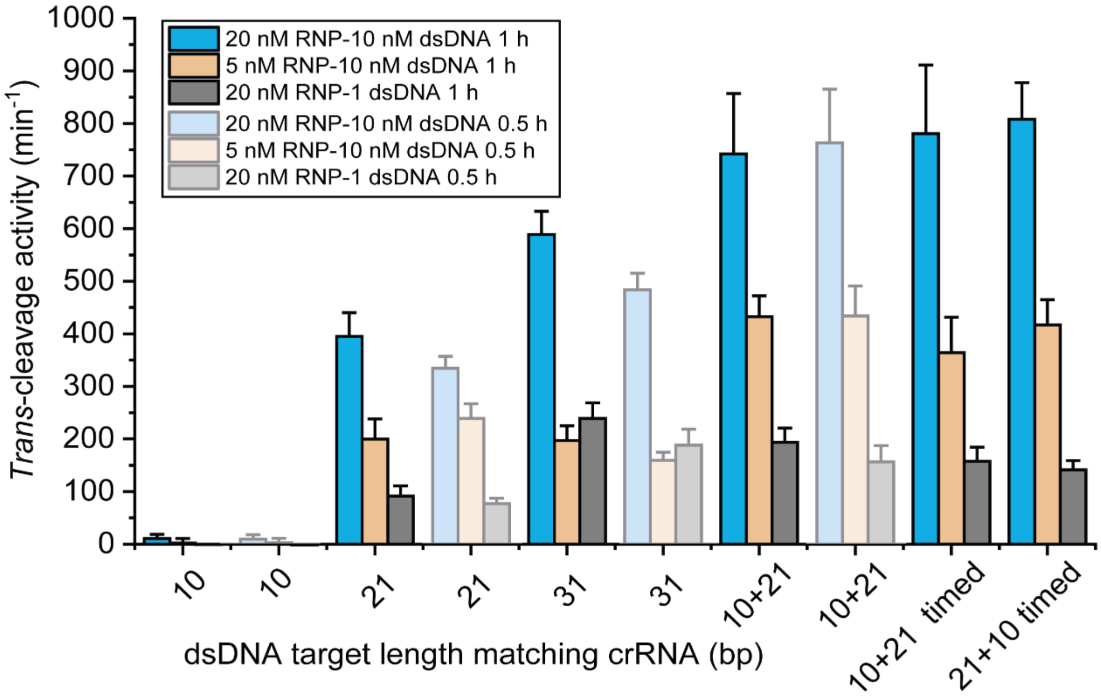


**Extended Data fig. 5, Absolute *trans*-cleavage rate of different fragment lengths for high GC-content MAL with 3, 21, 10+21 and 24 bp length crRNA matching fragments at different experimental conditions. Instead of in all other experiments, the DNA was first incubated with Cas12a for either 0.5 h or 1 h before the reporter fluorophore-quencher pair labelled DNA was added and the cleavage was followed as an increase in fluorescence signal. For the "timed" samples, the first fragment was added and incubated for 0.5 h before the second fragment was added and incubated for an additional 0.5 h (total incubation time 1 h). For 10nM dsDNA concentrations we observe a higher *trans*-cleavage for 3+21 compared to 24 nt, which is not in accordance with the other experiments where reporter DNA was immediately added to Cas12a. We hypothesize that this is possible due to the increased binding time, which allows more fragments to collectively bind to the crRNA. Experiments carried out with n=3 for the 0.5h incubation of the individual fragments and n=9 (three replicates for three independent targets) for all other experimental conditions.**

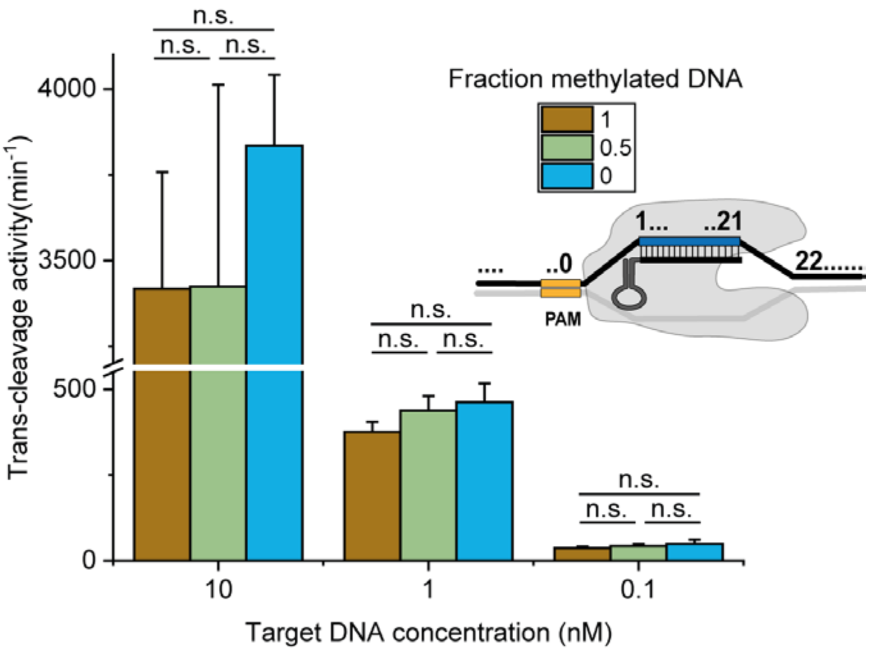


**Extended Data fig. 6, *trans*-cleavage rate of 0% Me, 50% Me and 100% Me MAL gene after mock digestion. Two-sample T test shows no significant differences in cleavage rate between the different methylation percentages. Experiments carried out with n=3.**

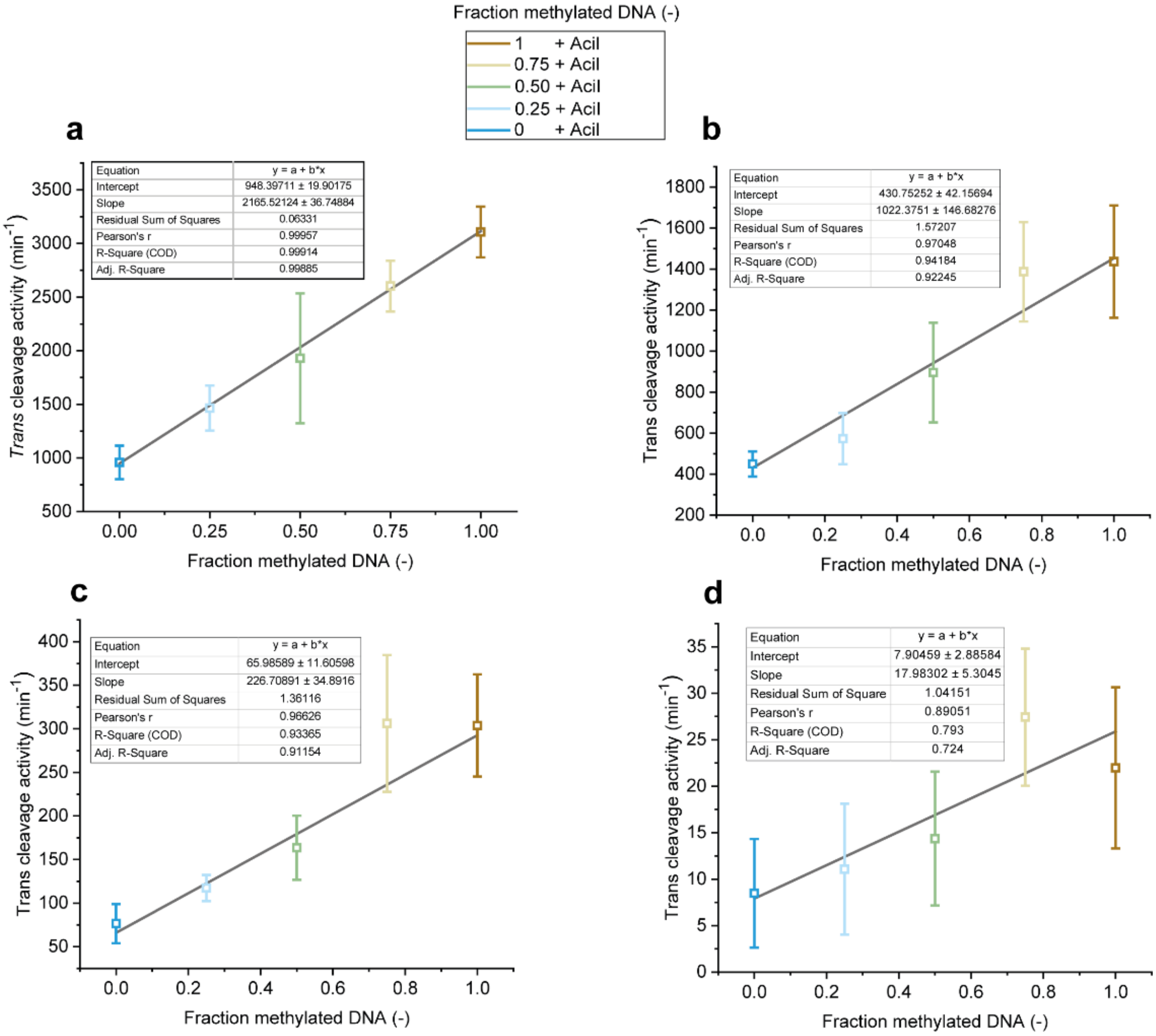


**Extended Data fig. 7, *Trans*-cleavage activity of different methylated DNA : non-methylated DNA fraction with 21 nt crRNA at a total DNA concentration of a) 10 nM. b) 5 nM, c) 1 nM, d) 0.1 nM. Error bars represent the mean +/- s.d., where n=9 (three replicates for three independent targets).**

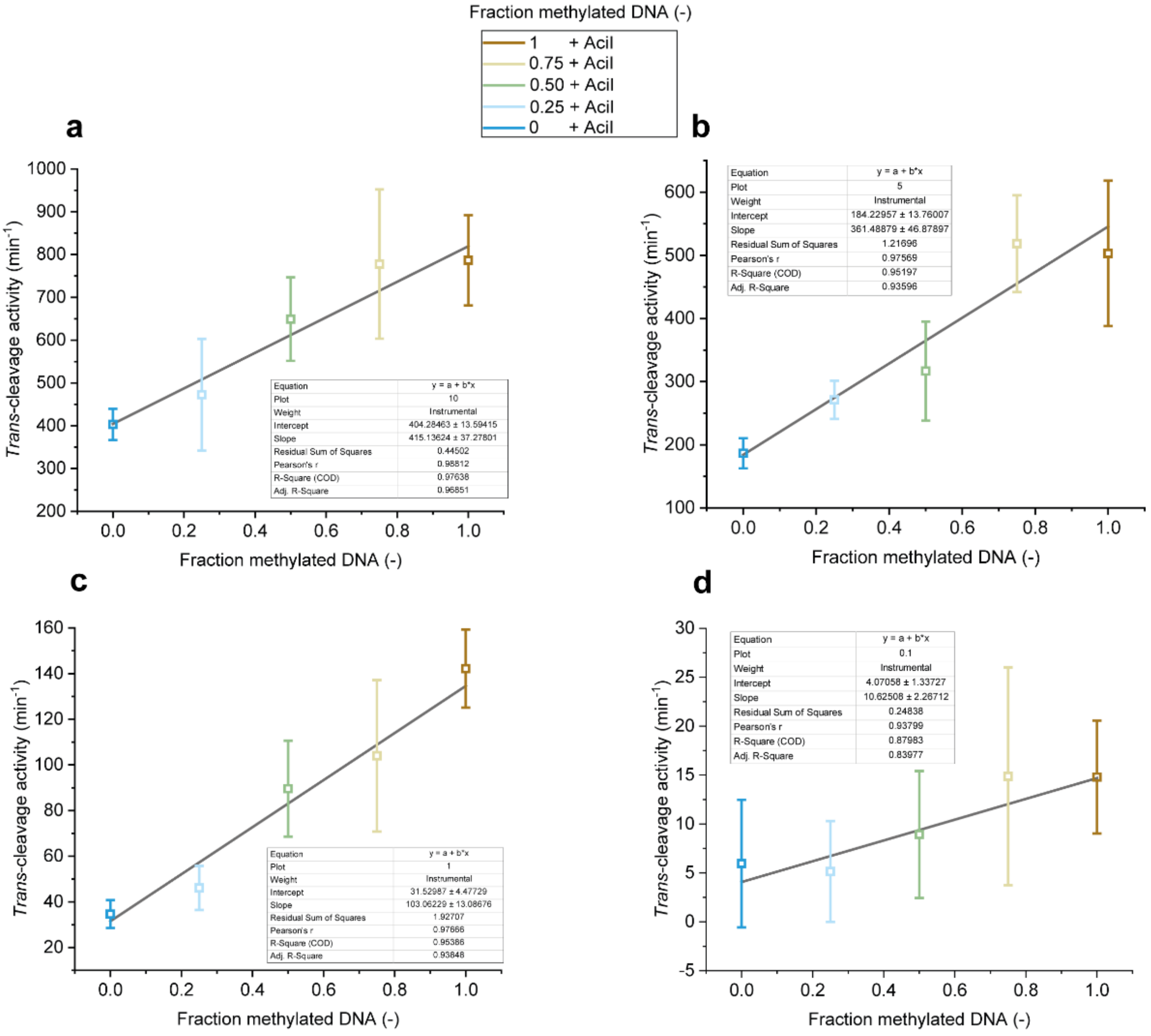


**Extended Data fig. 8, *Trans*-cleavage activity of different methylated DNA: non-methylated DNA fraction with 24 nt crRNA at a total DNA concentration of A) 10 nM. B) 5 nM, C) 1 nM, D) 0.1 n. Error bars represent the mean +/- s.d., where n=9 (three replicates for three independent targets).**
