## Supplementary material for "A CRISPR/Cas12a-assisted platform for identification and quantification of single CpG methylation sites": SI

*J.E. van Dongen[1]; J.T.W. Berendsen[1]; J.C.T. Eijkel[1]; L.I. Segerink[1]*
BIOS Lab on a Chip group, MESA+ Institute for Nanotechnology, Technical Medical Centre, Max Planck Institute for Complex Fluid Dynamics, University of Twente, Enschede, the Netherlands

### Supplementary information

#### Supplementary information 1: Mechanism behind Cas12a dsDNA sensing

Extended Data fig. 1 shows a schematic overview of the mechanism behind Cas12a's targeted *cis*- and non-targeted *trans*-cleavage. R-loop formation induces conformation activation of Cas12a, which is needed to reveal the RuvC catalytic site of the Cas12a effector protein (1C). This catalytic site is involved in the *cis-cleavage* of the non-target strand (light grey), the target strand (black) and the untargeted *trans*-cleavage of ssDNA. The R-loop formation, needed to induce this transition state, results in the formation of a (cr)RNA: DNA duplex. This duplex formation cannot be compared to the formation of RNA:DNA duplexes free in solution. In the presence of Cas12a more base pairs are formed than in regular RNA:DNA duplex formation where only a few base pairs contribute to the $k_d$ (binding affinity) and specificity. This late transition state results in a high specificity of Cas12a towards mismatches [1,2]. However, as discussed in the introduction, not all base pairs of the crRNA need to complement in order to achieve permanent binding of Cas12a. 15 out of 23 PAM-distal crRNA nucleotides complementing to the target sequence have shown to result in temporary binding and 17 out of 23 to permanent binding [3].

In a study performed by Swarts and Jinek, it was shown that *cis*-cleavage of the target DNA was needed, before *trans*-cleavage of the ssDNA could be initiated. By modifying the backbone of the dsDNA targets, the binding to Cas12a was still efficient, but due to the incomplete *cis*-cleavage, Cas12a's *trans*-cleavage activity was impaired [4]. From these results, the authors concluded that *cis*-cleavage is needed prior to *trans*-cleavage. On contrary, Chen et al. [5] showed that target strand cleavage by Cas12a is not required to trigger the *trans*-cleavage activity towards ssDNA. Short PAM containing target sequences, complementary to the crRNA of 10-25 bp in length, were incubated with Cas12a, and activity of the Cas12a was observed, while no *cis*-cleavage took place. From this we hypothesize that the most important prerequisite for *trans*-cleavage, after structural re-arrangement of Cas12a to reveal the RuvC catalytic site, is a clear RuvC site. This RuvC catalytic sited could be cleared either by *cis*-cleavage of target strands, resulting in diffusion of the PAM distal fragment or by short(er) target strands that do not interfere with this catalytic site (Figure 1D). Therefore, we hypothesize that *trans*-cleavage of shorter target DNA fragments is fully dependent on the R-loop formation step.

**Supplementary Information references:**

**Supplementary table 1: DNA sequences used in this study (5’-> 3’)**

**High GC-content MAL fragment sequences (5’->3’)**

| | |
|---|---|
| **MAL_10nt_FWD** | **GCG**GAAAAAT |
| **MAL_10nt_REV** | ATTTTTC**CGC** |
| **MAL_14nt_FWD** | **TCCAGCG**GAAAAAT |
| **MAL_14nt_REV** | ATTTTTC**CGCTGGA** |
| **MAL_18nt_FWD** | **CGCATCCAGCG**GAAAAAT |
| **MAL_18nt_REV** | ATTTTTC**CGCTGGATGCG** |
| **MAL_22nt_FWD** | **TTAACGCATCCAGCG**GAAAAAT |
| **MAL_22nt_REV** | ATTTTTC**CGCTGGATGCGTTAA** |
| **MAL_27nt_FWD** | **GCACTTAACGCATCCAGC**GGAAAAAT |
| **MAL_27nt_REV** | ATTTTTC**CGCTGGATGCGTTAAGTGC** |
| **MAL_31nt_FWD** | **TCGCGCACTTAACGCATCCAGCG**GAAAAAT |
| **MAL_31nt_REV** | ATTTTTC**CGCTGGATGCGTTAAGTGCGCGA** |
| **MAL_21nt_FWD** | **TCGCGCACTTAACGCATCCA** |
| **MAL_21nt_REV** | **TGGATGCGTTAAGTGCGCGA** |

**Low GC-content fragment sequences (5’->3’)**

| | |
|---|---|
| **LowGC_10nt_FWD** | GCTTTTC**TAC** |
| **LowGC _10nt_REV** | **GTA**GAAAAGC |
| **LowGC _14nt_FWD** | GCTTTTC**TACTTAA** |
| **LowGC _14nt_REV** | **TTAAGTA**GAAAAGC |
| **LowGC _18nt_FWD** | GCTTTTC**TACTTAAGCAT** |
| **LowGC _18nt_REV** | **ATGCTTAAGTA**GAAAAGC |
| **LowGC _22nt_FWD** | GCTTTTC**TACTTAAGCATTAGT** |
| **LowGC _22nt_REV** | **AATAATGCTTAAGTA**GAAAAGC |
| **LowGC _27nt_FWD** | GCTTTTC**TACTTAAGCATTAGTATAAT** |
| **LowGC _27nt_REV** | **TACTAATAATGCTTAAGTA**GAAAAGC |
| **LowGC _31nt_FWD** | GCTTTTC**TACTTAAGCATTAGTATAATGCAT** |
| **LowGC _31nt_REV** | **TACTAATAATGCTTAAGTA**GAAAAGC |

| | |
|---|---|
| **LowGC _21nt_FWD** | **TTAAGCATTAGTATAATGCAT** |
| **LowGC _21nt_REV** | **ATGCATTATACTAATGCTTAA** |

**Multiple restriction site sequence (5'-> 3')**

| | |
|---|---|
| **MRS_FWD** | ATTTTTC**CGGTATGCGCTAATGACGCGA**CAT |
| **MRS_REV** | ATG**TCGCGTCATTAGCGCATACCG**GAAAAAT |

**MAL 120 bp dsDNA fragment (5' -> 3')**

| | |
|---|---|
| **MAL_120nt_FWD** | CCCCAGCCTGTGGCGGTGGTCCAGTTCCGCCAGG<br>AAACCGCCGCCTGGAGCTGTGGG**TCGCGCACATT**<br>**AACGCATCCAGCG**GAAAAATGAAGGAGACCCAAA<br>TTCAAAGTTAAAGTAATG |
| **MAL_120nt_REV** | CATTACTTTAACTTTGAATTTGGGTCTCCTTCAT<br>TTTTC**CGCTGGATGCGTTAATGTGCGCGA**CCCAC<br>AGCTCCAGGCGGCGGTTTCCTGGCGGAACTGGAC<br>CACCGCCACAGGCTGGGG |
